# Life inside polymetallic nodules

**DOI:** 10.64898/2026.08.02.742282

**Authors:** Coral Diaz-Recio Lorenzo, Tanja Stratmann, Mark Eduard de Wilt, Ranju Radhakrishnan, Lara Macheriotou, Ellen Pape, Lucille Hoogerdijk, Nadinne Jeanneth van den Hooven-Iza, Jan Tobias Krüger, Lea Kaspersmeier, Moritz Neuser, Moritz Klöckner, Martin Lauth, Mirko Schaper, Tasnim Patel, Sabine Gollner

**Affiliations:** Department of Ocean Systems, NIOZ – Royal Netherlands Institute for Sea Research, Landsdiep 4, 1797 SZ ‘t Horntje (Texel), The Netherlands; Station Biologique de Roscoff (Sorbonne University), Pl. Georges Teissier, 29680 Roscoff, France; ERC Group “Benthic Ecological Biogeochemistry”, MARUM – Center for Marine Environmental Sciences, University of Bremen, Leobener Str. 8, Bremen 28359, Germany; Department of Earth Sciences, Faculty of Geosciences, Utrecht University, Vening Meineszgebouw A, Princetonlaan 8a, 3584 CB Utrecht, The Netherlands; IFREMER, Centre Bretagne, UMR BEEP, Univ, Brest, Z.I. de la Pointe du Diable, CS 10070, 29280 Plouzané, France; Marine Biology Research Group, Department of Biology, Ghent University, Krijgslaan 297, S8, 9000, Ghent, Belgium; Avans University of Applied Sciences, Breda (Biomedical Laboratory Research), Hogeschoollaan 1, 4818 CR Breda, The Netherlands; Chair of Materials Science, Paderborn University, Mersinweg 7, 33100 Paderborn, Germany; Marine Ecology and Management (MARECO), Royal Belgian Institute of Natural Sciences, Brussels, B-1000, Belgium

**Keywords:** GSR, BGR, Patania, deep-seabed mining

## Abstract

Deep-sea abyssal plains harbour extensive biodiversity, much of which remains undocumented. Polymetallic nodules in the abyssal plains of the Clarion-Clipperton Zone (CCZ), Central Pacific, harbour diverse meiofaunal communities within their crevices, representing an understudied habitat. Given their high concentrations of nickel, copper, and cobalt, nodules are currently one of the primary targets for the extraction of deep-sea mineral resources. Here, we characterise the physical properties of polymetallic nodules, quantify their meiofaunal crevice communities, and compare them to those from the nearby sediments. We show that nodules contain internal cracks and microfossils, that their crevices harbour meiofaunal communities dominated by nematodes and copepods, and that faunal abundance increases significantly with nodule volume. Nodule-crevice nematode and copepod abundances were equivalent to approximately 10% and 25%, respectively, of those recorded in nearby sediments across the two exploration contract areas in the eastern CCZ. Based on 18S rRNA data, nematode and copepod assemblages recovered from nodule crevices differed from those recovered from nearby sediments collected during the same expedition and within the same exploration contract areas. Our findings, therefore, point to the possibility of nodule-associated crevice meiofauna communities and the consequent risk of biodiversity loss associated with nodule removal through potential future commercial mining.

## Introduction

Deep-sea polymetallic nodules (potato-sized mineral accretions on the seabed; hereafter called nodules) were first discovered during the H.M.S. *Challenger* expedition in the late 19th century [1]. The Clarion-Clipperton Zone (CCZ), spanning the Pacific Ocean from Hawaii to Mexico (0°N, 160 W to 23.5°N, 111.4°W [2]; Fig. 1), hosts a high density of resource-grade nodules primarily composed of iron (Fe) and manganese (Mn) oxides, as well as potentially commercially valuable metals such as nickel (Ni), copper (Cu), and cobalt (Co) [3] . These metals, therefore, represent a major target for potential deep-sea mineral extraction. Nodules take millions of years to form [4,5], and organisms dependent on them will not recover post-mining as their habitat will be lost at the mined location [6,7]. Therefore, there is a need to characterise communities associated with nodule-crevices prior to resource exploitation contracts being granted by the International Seabed Authority (ISA), to obtain comprehensive baseline data that are critical for science-based environmental management.

**Figure 1.**
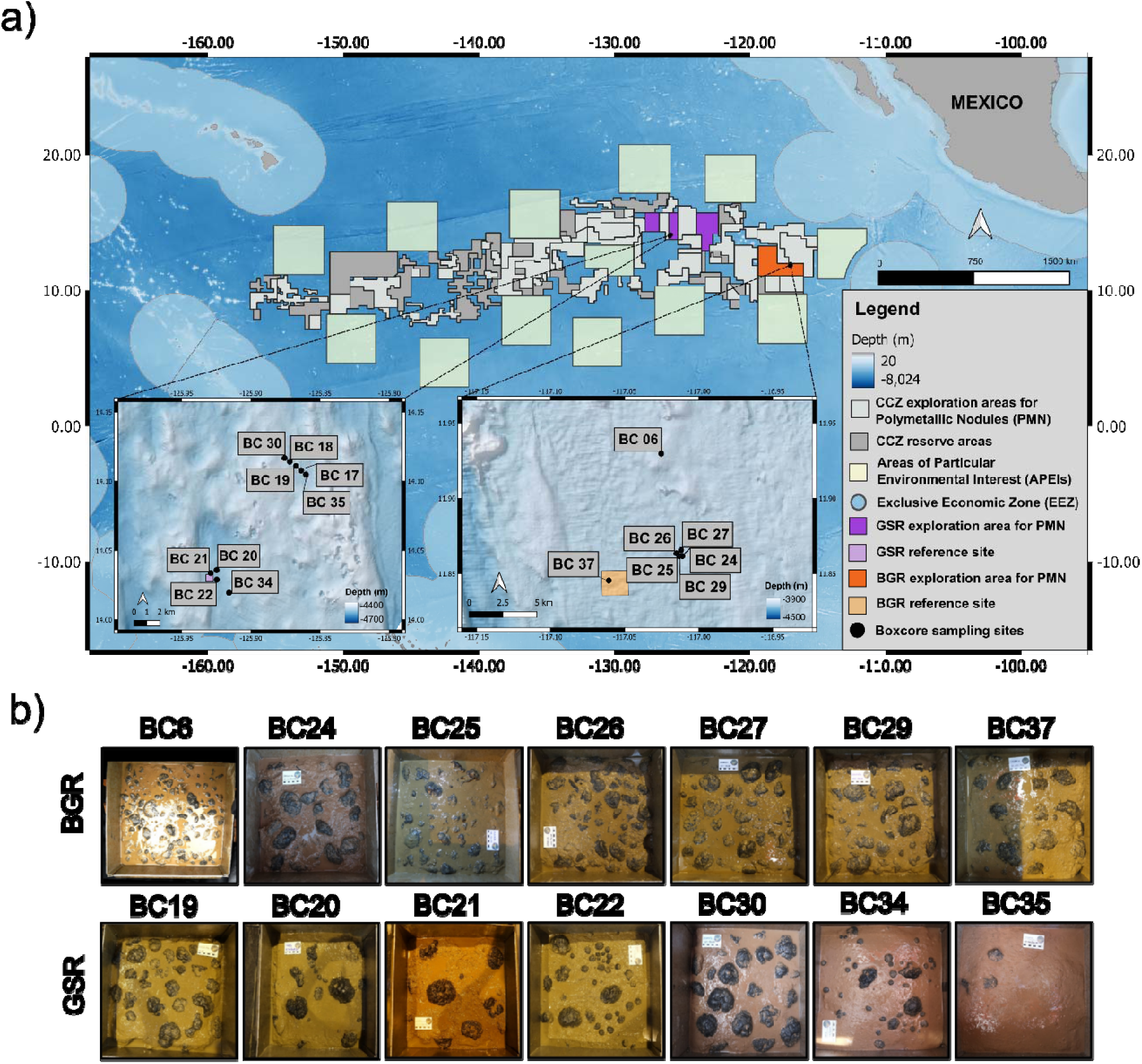
(a) Sampling locations in the Clarion Clipperton Zone highlighting the BGR and GSR exploration contract areas, and the box core sampling locations within each area during the SO268 expedition in 2019. Reference areas are nearby undisturbed seafloor sites used as baseline controls for comparison with the disturbed areas. Maps created in QGIS. (b) Photographs of the surface of each box core from which the nodules were collected (box core surface area = 2,500 cm²).

Over 5,000 undescribed benthic metazoan species have been recorded in the CCZ [8] and macrofaunal species richness curves show no signs of plateauing [9,10]. Numerous studies have examined biodiversity responses to benthic disturbance [6,11–18], showing that mining simulation conducted with non-mining equipment and test mining conducted with (pre-)prototype collector vehicles cause immediate losses in faunal abundance and diversity within the path of direct disturbance. While these studies show alterations in the sediment fauna, there is an urgent need for baseline data on nodule-crevice fauna as these would be directly removed by mining.

To date, only seven studies have investigated nodule-crevice fauna, all based on morphology. Initial reports of nodule-crevice fauna described a new tardigrade species from a nodule in the Peru Basin (Southeast Pacific) [19]. Subsequent studies reported marked differences in nematode genus composition between nodule crevices and surrounding sediments [20,21], followed by the first account of nodule-crevice foraminifera, which, while not meiofauna, are found in the same size fraction (32µm – 1mm) [22]. Later work in the Indian Ocean showed that nematode communities differed mainly in the relative abundance of shared genera between substrates, while forming distinct assemblages at the species level [23]. A similar pattern was reported from the Global Sea Mineral Resources (GSR) mining exploration contract area in the CCZ, where community differences were associated with nematode buccal morphology, indicating contrasting feeding strategies between sediment and nodule assemblages [24]. These studies have focused on Nematoda, while Copepoda have never been investigated quantitatively and molecular studies on nodule-crevice fauna are, to date, lacking.

Given the need for more baseline data on this particularly vulnerable group, our study employed a multifaceted approach to understand nodules as habitats for nodule-crevice fauna. We used scanning electron microscopy (SEM), Energy Dispersive X-ray spectroscopy (EDX), and digital microscopy (DM) to characterise nodule porosity and crack diameter, and to map the elemental composition of cracks of 24 nodules collected from the Bundesanstalt für Geowissenschaften und Rohstoffe (BGR, Germany) and Global Sea mineral Resources (GSR, Belgium) nodule exploration contract areas of the CCZ. In addition, we quantified the nodule-crevice fauna from 244 nodules and investigated the relationship between meiofauna abundance, size, and biomass in relation to the volume of those 244 nodules (Nodule dimensions and sizes can be found in Schoening & Gazis 2019 [25]. Finally, we used DNA barcoding and metabarcoding to investigate the phylogeny and community structure of the crevice fauna and compared it to the community structure from nearby sediments in the same locations.

## Results and discussion

### Physical properties and geochemical composition of nodules

DM and SEM revealed bright and dark layers arranged in massive, laminated and dendritic structures (Fig. 2a, Supplementary Fig. 1). Crevice-count relationships with ellipsoid volume differed among nodule sections (χ²(2) = 8.52, *n*_nod_ = 24, p-value = 0.014). Counts were unrelated to volume in central and top sections, but decreased by 14.8% for each twofold increase in volume in bottom sections (z = -2.05, 95% confidence intervals (CI) = 0.7 – 26.9%, *n*_nod_ = 7, p-value = 0.041). Porosity averaged approximately 23% and was unrelated to ellipsoid volume (z = -0.36, *n*_nod_ = 24, p-value = 0.721), although it was higher in the GSR than in the BGR (z = 3.03, odds ratio (OR) = 1.44, 95% CI = 1.14 – 1.83, *n*_nod_ = 24, p-value = 0.002). Median crevice area showed a concave, unimodal relationship with ellipsoid volume (χ²(1) = 7.70, *n*_nod_ = 24, p-value = 0.0055), with a fitted maximum at approximately 92 ml. For each twofold increase in volume, internal and external crack diameters increased by 15.7% (z = 6.26, 95% CI =10.6 – 21.2%, *n*_nod_ = 24, p-value < 0.001) and 15.4% (z = 3.93, 95% CI = 7.4 – 23.9%, *n*_nod_ = 24, p-value <0.001), respectively. External crack diameters were also 32% greater in GSR than in BGR (z = 2.14, 95% CI = 1.02 – 1.71, *n*_nod_ = 24, p-value = 0.033). Thus, larger nodules contained wider crevices without a corresponding increase in total porosity (Supplementary Fig. 2, 3).

**Figure 2.**
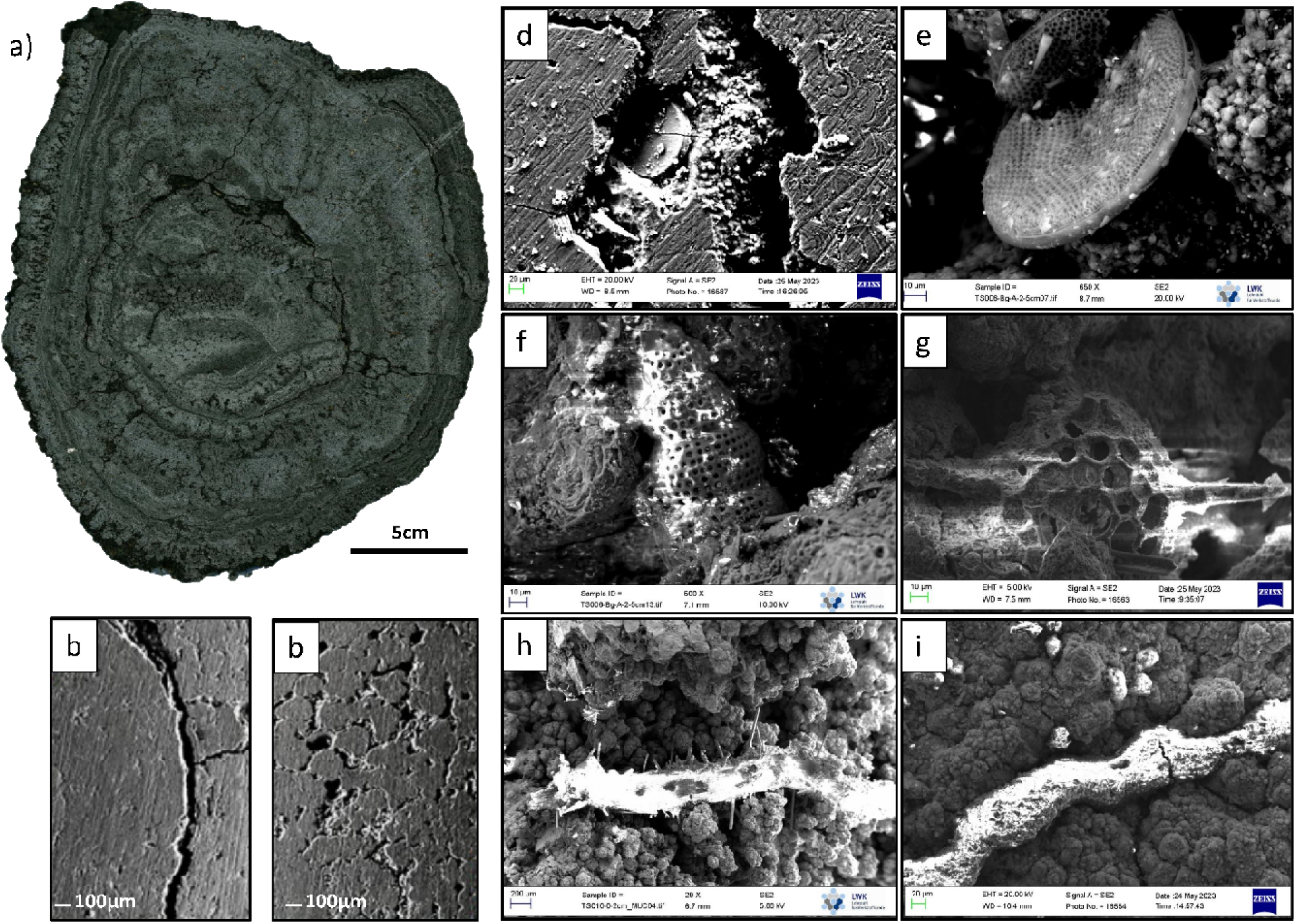
Example of a polished polymetallic nodule cross section and examples of microfossils that can be found inside a nodule. (a) Central cross section of a very small nodule (0 – 10 ml volume), (b) example of a cross section along a crack from a polished nodule section, (c) example of multiple nodule cracks and pores, (d) coccoid microorganisms growing on a *Coscinodiscus*-type diatom, (e) centric diatom likely *Kozloviella subrotunda*, (f) Nasselarian radiolarian likely *Stichocorys* sp., (g) Spumellarian radiolarian, (h) agglutinated Foraminifera identified as *Saccorhiza ramose*, (i) agglutinated Foraminifera identified as *Tolypammina* sp.

Maps of SEM-EDX showed relatively even distributions of oxygen (O), while Mn, Fe, chloride (Cl) and aluminium (Al) were structured in clear layers (Supplementary Fig. 4). Layer compositions differed between contract areas: layers of nodules from the BGR area showed lower Mn/Fe-ratios (6.38) and higher (Ni+Cu)/Co-ratios (39.0) than those from GSR (Mn/Fe-ratio: 7.12, (Ni+Cu)/Co-ratio: 30.1). Ternary classifications indicated that nodules from both areas contained layers formed via hydrogenetic growth (layer type 1), diagenetic growth (types 2.1, 2.2) and a mixed type (layer 3); Supplementary Fig. 5), consistent with established models for nodules [26–29]. Hydrogenetic layers accreted slowly, at only a few mm My^-1^, whereas diagenetic layers accreted 10s to 100s mm My^-1^, with mixed layers showing intermediate rates. These alternations point to repeated switches between hydrogenetic and diagenetic growth modes through time, consistent with established models of nodule growth.

Interpreted mechanistically, type 1 (hydrogenetic) layers reflect oxic bottom-water or shallow-pore-water precipitation during phases of stronger deep-water ventilation (e.g., vigorous Antarctic Bottom Water [29]) whereas diagenetic layer types 2.1 – 2.2 reflect suboxic upper-sediment pore waters associated with higher organic-carbon export and/or weaker ventilation [28]. Further, pores/cracks in the different nodule areas (Fig. 2b, c; nodule surface vs nodule centre) were connected, sometimes even via cracks to the nodule surface. Such networks may permit oxygen and nutrient transport into nodule interiors and support manganese- and iron-cycling microorganisms, as described by Blöthe et al. (2015) [30].

The walls of the cracks inside the nodules had lower Mn/Fe ratios (3.65) than dense rings (compact, metal-rich accretion layers) (8.82), meaning that crack walls were relatively enriched in Fe and depleted in Mn. Dense rings, in turn, reflected primary accretion fabrics with higher Mn and Co + Ni + Cu, characteristic of compact growth laminae.

Blöthe et al. (2015) [30] also reported lower Mn/Fe ratios in pore-filling material than in adjacent dendritic structures from GSR nodules and showed that interconnected pores and cracks enable oxygen and nutrient transport into the nodule interior. They identified Gammaproteobacteria, mainly *Shewanella* and *Colwellia*, associated with Mn cycling, and predominantly coccoid cells ∼1 µm in diameter. Our SEM images similarly revealed coccoid microorganisms (∼1 – 4 µm) on internal mineral surfaces and within cracks (Fig. 2d), comparable to those reported from other CCZ nodule crevices [31]. Yapan et al. (2026) [31] recently showed that nodule crevice bacterial communities varied strongly among nodule parts (top, bottom and central sections), consistent with pronounced internal microbial heterogeneity that may accompany fine-scale differences in metal composition within nodules. However, without 16S rRNA gene sequencing or functional assays, the observed microorganisms in this study cannot be assigned taxonomy or metabolic function. Conversely, as Yapan et al. (2026) [31] did not measure mineralogy, this link is theoretical. Lower Mn/Fe ratios along crack walls may therefore reflect secondary mineral precipitation, abiotic alteration, microbial metal transformations or their interaction. The co-occurrence of coccoid microorganisms and contrasting Mn/Fe signatures is consistent with, but does not demonstrate, a microbial contribution to fine-scale geochemical heterogeneity within nodules.

### Microfossil assemblages

Microfossils are widely reported from nodules [32–34]. In this study, SEM imaging revealed abundant filaments, sponge spicules, diatoms (centric and pennate forms), radiolarians (order Nassellaria and Spumellaria) and agglutinated Foraminifera embedded within the nodule matrix and along crack walls (Fig. 2d – i). Foraminifera, diatoms, and radiolarians showed similarly variable elemental signatures within and between nodules [35]. Elemental signals with mixed silicon (Si), Al, and sodium (Na) may reflect agglutinated wall material that Foraminifera might have incorporated locally from fine sediment. Agglutinated Foraminifera such as *Rhizammina* sp., *Tolypammina* sp. and *Saccorhiza ramosa* identified in our SEM images, most likely represented crevice-dwelling colonisers that were subsequently preserved within the nodules. In contrast, diatoms and radiolarians may have either settled onto the nodule surface during primary growth and become overgrown or been introduced later through cracks during cementation [36]. Consequently, these siliceous microfossils do not provide a direct age for the initiation of nodule growth. The stratigraphic ranges of taxa such as the diatom *Kozloviella subrotunda* and the radiolarian *Stichocorys* spp. may constrain the age of material incorporated into nodule layers. However, these microfossils may have been reworked from older sediments and subsequently incorporated into younger nodule layers. Their stratigraphic ages therefore do not necessarily represent the timing of mineral-layer formation or the initiation of nodule growth.

### Quantitative analysis of nodule-crevice organisms

From 244 processed nodules collected in the BGR (*n*_nod_ = 153) and GSR (*n*_nod_ = 91), we extracted a total of 4,139 individuals from nodule-crevices, both from nodules fixed in formaldehyde and ethanol [37]. Communities were composed of Nematoda (52.9 – 84.0% relative individual abundance), followed by Copepoda (7.41 – 13.4%), Foraminifera (5.64 – 29.4%) and polychaetes (2.24 – 2.47%) (Fig. 3, Supplementary Table 1), consistent with previous reports [21,24,38]. In our study, the mean number of individuals per nodule was 17, lower than that reported by Bussau et al. 1995 [38] (116 ind. nodule^-1^) and Pape et al. 2021 [24] (28 ind. nodule^-1^) while a maximum of 245 individuals in a single nodule (this study) exceeded previously reported maxima (170 ind. nodule ¹ and 86 ind. nodule ¹, respectively [21,24]), reflecting the broader nodule size range analysed here.

**Figure 3.**
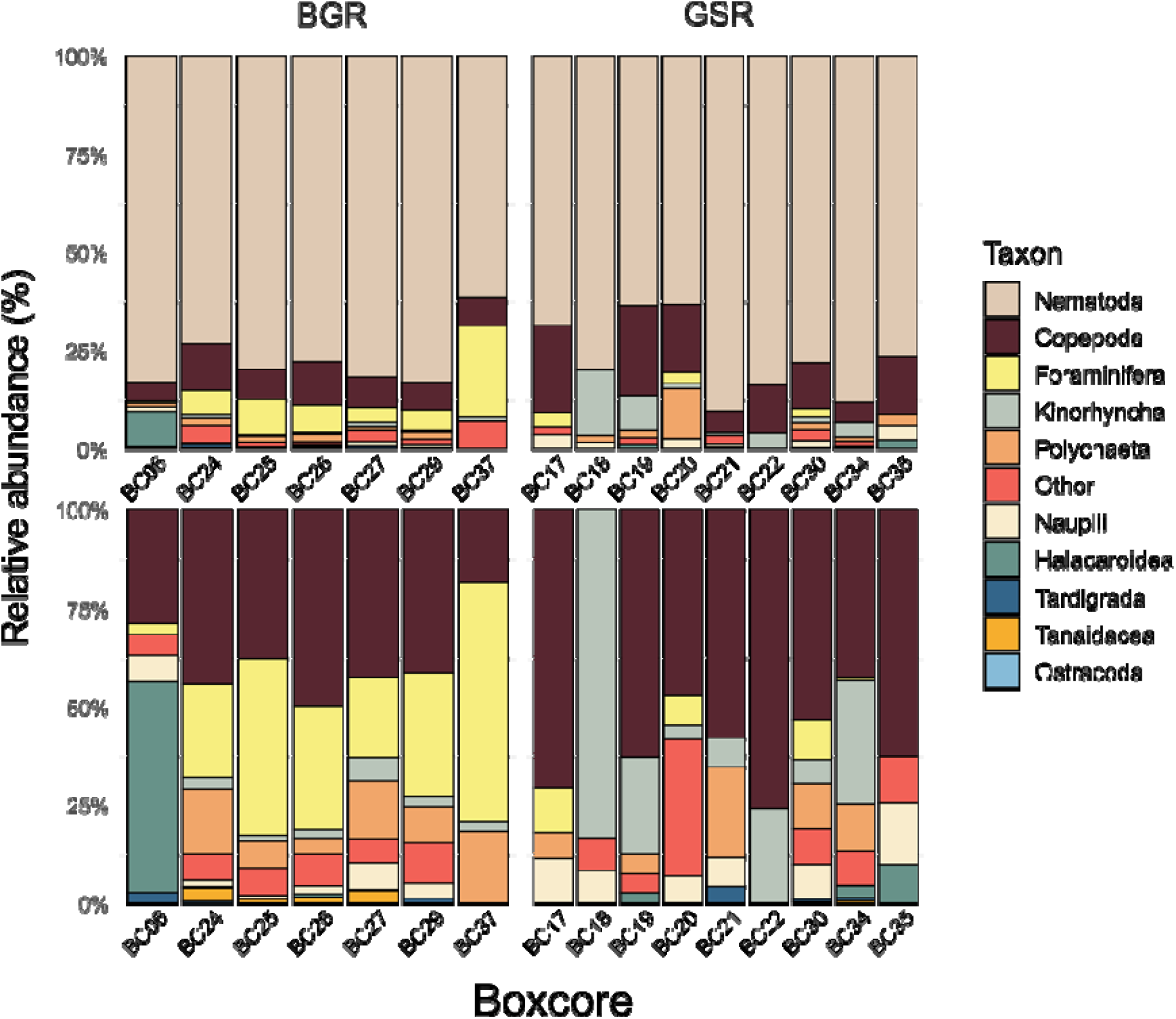
Higher taxon composition of nodule-crevice organisms in both the BGR and GSR contract areas, including (top panels) and excluding (bottom panels) Nematoda for better visualisation of rare taxa. “Other” refers to unidentified biological material. Legend is ordered by relative abundance.

Nodule-crevice Nematoda densities (in ind. 10 cm ² of nodule surface area, i.e., the amount of nodule area covering the seabed) in the BGR and GSR ranged from 1.27 to 8.08 (5.35±2.40, *n*_nod_ = 153) and from 2.10 to 26.8 (7.52±7.60, *n*_nod_ = 91), respectively, whereas nodule-crevice Copepoda densities ranged from 0.21 to 1.49 (0.70±0.42, *n*_nod_ = 153) and from 0.00 to 5.50 (1.30±1.63, *n*_nod_ = 91), respectively. In contrast, sediment Nematoda densities (ind. 10 cm ² of sediment surface area) in the BGR and GSR ranged from 44.6 to 132 (88.3±61.7, *n* = 2) and from 6.36 to 136 (61.5±33.8, *n* = 14), respectively, while sediment Copepoda densities ranged from 2.49 to 4.01 (3.25±1.07, *n* = 2) and from 0.83 to 8.01 ind. 10 cm ² (4.33±2.34, *n* = 14), respectively [39,40]. These results support the findings from Singh et al. 2019 [23] and Pape et al. 2021 [24] that nodules host fewer individuals per unit area than in nearby sediments and also that strong heterogeneity exists among nodules as within sediments, a prominent feature of the CCZ benthos. Compared with abundances reported from sediments sampled during SO268-1/2 in the BGR [40] and GSR exploration contract areas [39], nodule crevices supported nematode and copepod abundances equivalent to ∼10% and ∼25% of sediment values, respectively, indicating a non-negligible contribution to local meiofaunal standing stocks. These comparisons should be interpreted cautiously because nodule-crevice abundances in this study were derived from box cores in the BGR and GSR, whereas sediment abundances were derived from multicores in the same location. Sediment abundances were restricted to the upper 0 – 3 cm and standardised to 10 cm², minimising differences in sampled depth and area between box core and multicorer samples. Although minor gear-related differences in recovery and disturbance of the sediment surface may remain, the datasets provide a reasonable first-order comparison of faunal density.

Nodule-crevice faunal abundance increased significantly with nodule volume. Each twofold increase in volume was associated with increases of 34.4% in total abundance (95% CI: 27.5 – 41.6%, *n*_observation_ = 244, p-value <0.001), 35.7% in Nematoda, 37.4% in Copepoda, 26.5% in Foraminifera, and 65.8% in Polychaeta (Supplementary Table 2). Although larger nodules contained more fauna overall, abundance did not increase in direct proportion to nodule volume. Consequently, smaller nodules supported more individuals per unit volume. Combined nematode and copepod biomass increased by 32.8% per doubling of volume (95% CI: 16.9 – 50.9%, *n*_observation_ = 57, p-value <0.001), whereas nematode length (two-fold increase: -0.2%, 95% CI: - 2.3 – 1.9%, *n*_observation_ = 875, p-value = 0.822) and copepod width (change per two-fold increase: +3.4%; 95% CI: -0.1 – 7.0%, *n*_observation_ = 882, p-value = 0.057) did not vary significantly. At equivalent nodule volumes, total abundance, and nematode and copepod abundances were respectively 72.5% (95% CI: 3.5 – 188%, *n*_observation_ = 244, p-value = 0.040), 108% (95% CI: 14.0 – 279%, *n*_observation_ = 244, p-value = 0.021), and 121.5% (95% CI: 8.2 – 353%, *n*_observation_ = 244, p-value = 0.035) higher in the GSR than in the, whereas foraminiferal abundance was 83.4% lower in the GSR (95% CI: -92.1 – -65.3%, *n*_observation_ = 244, p-value = <0.001). Polychaete abundance did not differ significantly between regions (higher abundance in GSR contract are compared to BGR contract area: -21.7%; 95% CI: -57.1 – 42.8%, *n*_observation_ = 244, p-value = 0.420). Analyses were tested with equal number of nodules per size class confirming that the higher proportion of small nodules did not affect the positive relationship between nodule volume and faunal abundance (Supplementary Fig. 6, 7).

### Nematoda crevice community

Nematode communities from 17 nodules (each nodule was treated as one sample) collected from six box cores were investigated using metabarcoding (Supplementary Data 1). A total of 67 amplicon sequence variants (ASVs) were recovered, from which Bayesian Poisson Tree Processes (bPTP) [39] delimited 22 putative species (Supplementary Data 2). Per-nodule richness and Shannon diversity were low (S = 1 – 6, H′ = 0 – 1.67), whereas Pielou’s evenness was high (J′ = 0.94 – 1.00; Supplementary Table 3).

To avoid pseudo replication, ASV occurrences were aggregated by independent sampling unit: box cores for nodules and multicores for sediments. Nodule nematode ASV composition did not differ detectably between the BGR and GSR contract areas (Permutational multivariate analysis of variance PERMANOVA: R² = 0.19, F(1,4) = 0.91, p-value = 1.00), and multivariate dispersion was comparable (PERMDISP: F(1,4) = 0.28, p-value = 0.90). Within sediments, contract area explained 10.3% of compositional variation (PERMANOVA: R² = 0.10, F(1,14) = 1.61, p-value = 0.01). However, sediment assemblages showed marked heterogeneity among multicore samples within contract areas, and the extent of this variability differed between the BGR and GSR (PERMDISP: F(1,14) = 78.7, p-value <0.001).

Nematode ASV composition differed significantly between nodule and sediment habitats. After accounting for exploration contract area, habitat explained 11.0% of compositional variation (PERMANOVA: R² = 0.11, F(1,19) = 2.56, p-value <0.001), whereas exploration contract area explained 5.6% (R² = 0.06, F(1,19) = 1.31, p-value = 0.04). Multivariate dispersion did not differ significantly between habitats (PERMDISP: F(1,20) = 3.57, p-value = 0.07), supporting a consistent habitat-related shift in ASV composition rather than an effect driven by differences in within-habitat variability.

Similarity percentage (SIMPER) analysis [41] identified *Acantholaimus* sp. and *Desmoscolex* sp. as the principal contributors to habitat dissimilarity, accounting for 18.3% and 14.1%, respectively. *Halalaimus* sp. contributed 5.9%, while *Chromadorita* sp. and *Phanodermopsis* sp. contributed 2.1% and 1.3%, respectively. Together, these five genera accounted for 41.7% of average dissimilarity (Supplementary Table 4). *Acantholaimus* sp. was frequently detected in sediments [16,17], but it was represented by only one ASV in nodules, whereas *Metoncholaimus* sp. and *Onchium* sp. occurred exclusively in nodule crevices. *Phanodermopsis* sp., *Oxystomina* sp., *Chromadorita* sp., and *Deontolaimus* sp. were mainly associated with nodule crevices, whereas *Desmoscolex* sp., *Manganonema* sp., and *Leptolaimus* sp. occurred only in sediments (Supplementary Data 3, Fig. 4), despite previous morphological records from GSR nodules [24].

**Figure 4.**
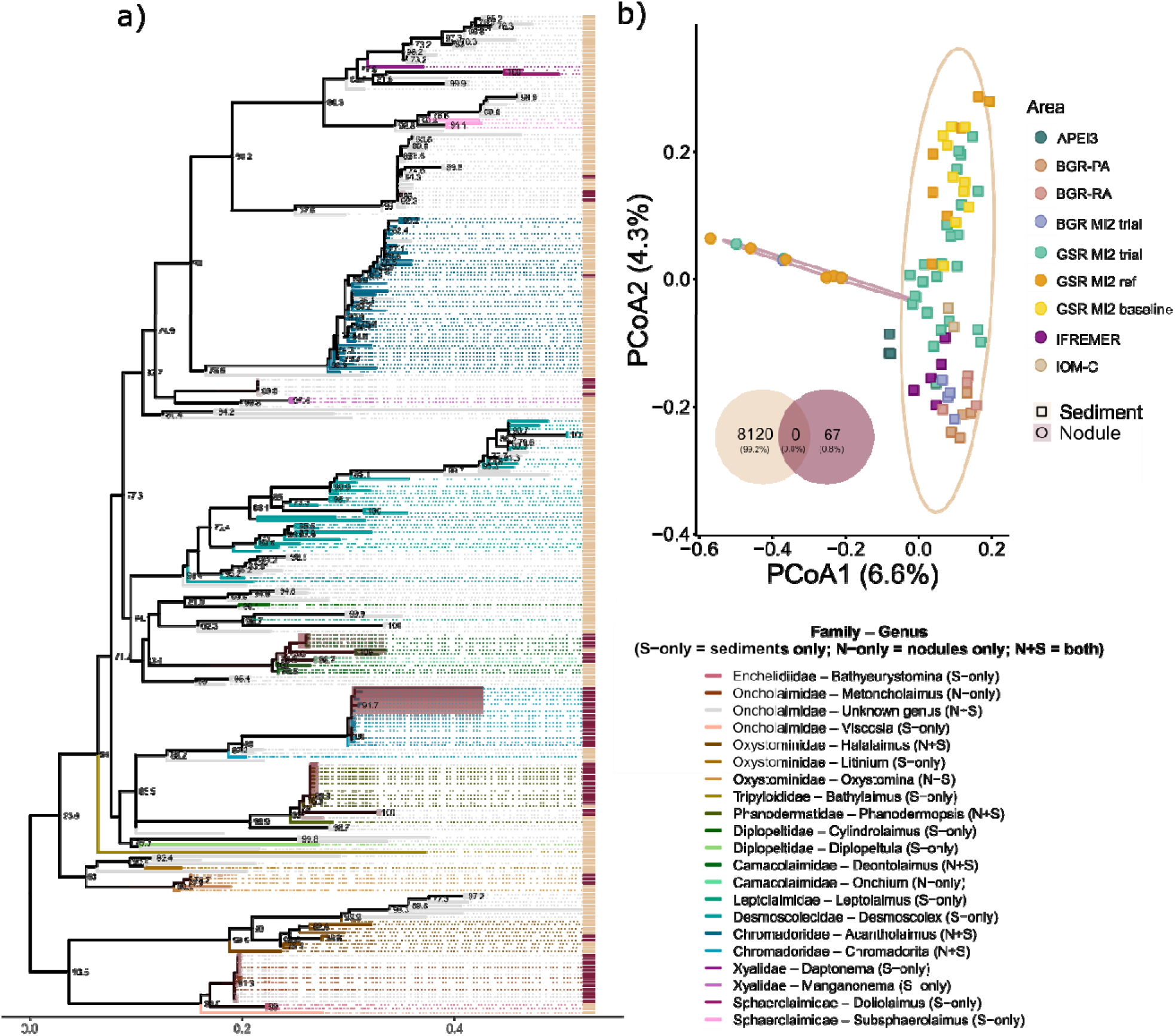
(a) Maximum likelihood phylogenetic construction for the Nematoda ASVs, where branches are coloured by Family-genus and correspond to either sediment or nodule samples. Shaded clades represent clades represented by >50% nodule samples. Values on the branches represent bootstrap values >70% indicative of well-supported diverging lineages. (b) Principal coordination analysis (PcoA) based on Sørensen distance (presence-absence) and number of shared sequences between nodules and sediments for Nematoda.

### Qualitative analysis of Copepoda

Copepoda from nodule crevices, 107 copepod individuals were successfully barcoded, from which Bayesian Poisson Tree Processes (bPTP) delimited 83 putative species. Across nodules, richness ranged from 1 to 11 species, Shannon diversity from 0 to 2.4 and Pielou’s evenness from 0.72 to 1.00 (Supplementary Table 5).

Sequence occurrences were once again grouped by independent sampling unit as for the nematodes (i.e., box core for nodules and multicores for sediments). Nodule copepod community composition did not differ significantly between the BGR and GSR exploration contract areas (PERMANOVA: R² = 0.108, F(1,11) = 1.327, p-value = 0.156), and multivariate dispersion was comparable (PERMDISP: F(1,11) < 0.001, p-value = 0.976). Differences between exploration contract areas for sediment communities could not be tested because no independent BGR sediment multicores were available.

Copepoda community composition differed significantly between nodule and sediment habitats (Fig. 5). After accounting for contract area, habitat explained 13.2% of compositional variation (PERMANOVA: R² = 0.132, F(1,20) = 3.247, p-value = 0.0001), whereas exploration contract area explained 4.7% and was not significant (R² = 0.047, F(1,20) = 1.148, p-value = 0.234). The comparison comprised 13 nodule box cores and 10 sediment multicores. As sediment units were available only from the GSR, the habitat effect was supported principally by the contrast between nodule and sediment assemblages within the GSR. Multivariate dispersion did not differ significantly between habitats (PERMDISP: F(1,21) = 4.034, p-value = 0.052), supporting a habitat-related shift in community composition, although some contribution from differences in within-habitat variability cannot be excluded.

**Figure 5.**
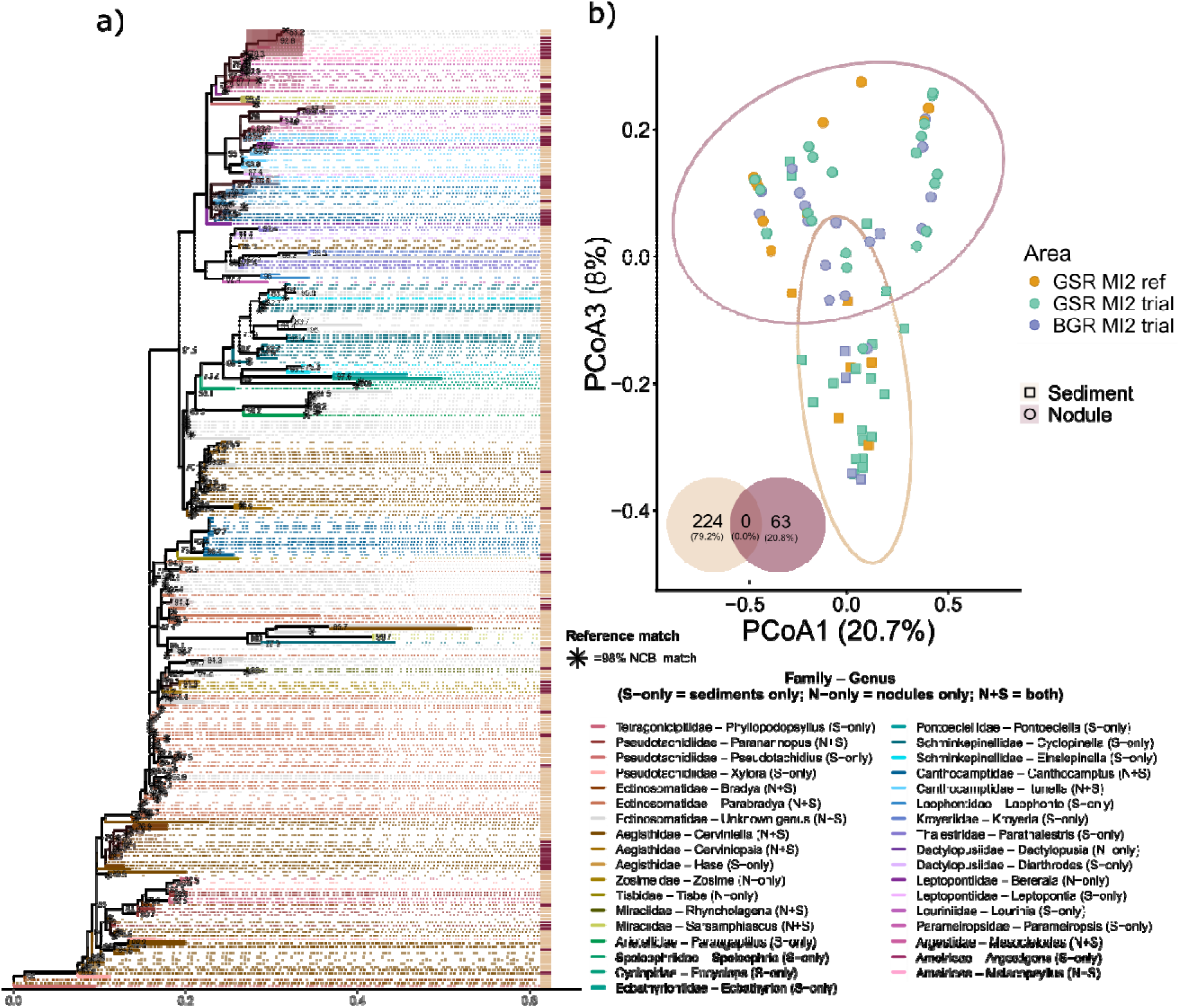
(a) Maximum likelihood phylogenetic construction for the Copepoda, where branches are coloured by Family-genus and correspond to either sediment or nodule samples. Stars represent sequences that matched with a reference sequence in NCBI with a percentage similarity of 98% or above, representing a “species” match. Values on the branches represent bootstrap values > 70% indicative of well-supported diverging lineages. (b) PcoA based on unweighted UniFrac distance (i.e., a phylogeny-informed presence–absence distance [46]), and number of shared sequences between nodules and sediments for Copepoda.

SIMPER analysis was restricted to the GSR, comprising nodules from eight box cores and sediment samples from ten multicores. *Cerviniella* sp. contributed most strongly to the average habitat dissimilarity, accounting for 25.5%, followed by *Parabradya* (13.4%), *Canthocamptus* (10.9%), *Itunella* (5.0%) and *Cyclopinella* (4.0%). Together, these five identified genera accounted for 58.8% of the average dissimilarity between nodule and sediment copepod assemblages (Supplementary Table 6). Sequences that differed from NCBI reference sequences by >2%, were treated as distinct molecular lineages (putative taxa), potentially reflecting undescribed diversity, but true molecular species boundaries require confirmation with higher-resolution markers (e.g., COI/28S) complemented by morphology. Phylogenetic data showed that the Canthocamptidae family was predominantly found in nodules, while other families like Gisilenidae and Cyclopidae were exclusive to sediments, however these results are based on NCBI-assigned taxonomy, and morphological taxonomic identification, would be needed to corroborate these results (Supplementary Data 4). The lack of shared sequences between nodules and sediments may be due to differences in feeding type or nutritional sources as has been suggested for Nematoda. Copepod community composition is known to respond strongly to habitat type, the amount and type of food available [42,43], and bacterial communities differ markedly between nodules and surrounding sediments [31,44,45], likely promoting differences in nutritional source. Together, these factors likely reinforce the observed segregation between nodule interiors and nearby sediments.

### Qualitative analysis of Foraminifera

We morphologically identified 67 foraminiferal individuals from nodule-crevices, comprising six species and 14 morphotypes (Supplementary Table 7). Foraminifera from the CCZ sediments are known for their remarkable diversity [47,48], making the results of this study particularly noteworthy as nodule-crevices were dominated by one species, namely *Thurammina* sp. 1. However, identification of Foraminifera, particularly soft-bodied monothalamids, is hindered by high morphological diversity, increasing the likelihood of misidentification and exclusion, thus diversity reported here may be underestimated. Of the six nodule-crevice species, three belonged to the genus *Thurammina* (*Thurammina.* sp. 1, *Thurammina* sp. 2, *Thurammina* sp. 3), one to *Textularia* (*Textularia* sp.), and two to organic-walled allogromiids (*Placopsinella-like* sp., *Hospitella* sp.). We also identified horseshoe-like, elongated forms and spheres located within empty radiolarian tests (Supplementary Table 7, Supplementary Fig. 8).

Foraminiferal morphotype community composition was examined among 32 nodules containing at least one recorded morphotype, representing 67 records across 14 taxonomic or morphological categories and eight box cores. Composition did not vary with nodule volume after accounting for box core structure and restricting permutations within box cores (PERMANOVA: R² = 0.019, pseudo-F(1,23) = 0.550, p-value = 0.878). This result was unchanged using presence-absence and Sørensen dissimilarity (R² = 0.014, pseudo-F(1,23) = 0.408, p-value = 0.930), or when box cores represented by only one nodule were excluded. At the box core level, pooled morphotype composition did not differ between BGR (*n*_box cores_ = 5) and GSR (*n*_box cores_ = 3) (PERMANOVA: R² = 0.133, pseudo-F(1,6) = 0.918, p-value = 0.414), and multivariate dispersion was homogeneous between regions (PERMDISP: F(1,6) = 1.357, p-value = 0.357). The regional result was likewise unchanged using presence-absence and Sørensen dissimilarity (PERMANOVA: R² = 0.086, pseudo-F(1,6) = 0.566, p-value = 0.808; PERMDISP: F(1,6) = 0.246, p-value = 0.736).

Unlike rigid calcareous forms, agglutinated and organic-walled taxa, can abandon and rebuild tests or infiltrate ultrafine fissures [22]. Earlier work has documented nodule-encrusting (*Tolypammina* sp.) or nodule-crevice-dwelling (*Saccorhiza ramosa*) Foraminifera, emphasizing surface and internal nodule-crevices as refugia and resource-rich microhabitats [49–51]. The reddish, Fe-rich agglutinated tests of the dominant species *Thurammina* sp. 1, suggest an affinity with bioavailability of iron oxides [52], while the consistent occurrence of *Thurammina* sp. 1 in cracks and high abundance in nodule-crevices relative to sediments implies adaptation to sheltered, particle-retaining spaces. Further, its capacity to vacate and reconstruct tests, likely facilitates the colonisation of irregular micro-spaces [53] such as nodule-crevices, while organic-walled species appear to exploit fine-sediment and/or detritus trapped in crevices. These results point to species such as *Thurammina* sp. 1 as a characteristic taxon for nodule communities [22,47] that would be particularly at risk should mining of deep-sea nodules occur.

### Nodules as habitats

Our analyses demonstrate that nodule crevices function as distinct, three-dimensional habitats that harbour different benthic communities from those in nearby sediments. Nodule architecture, geochemistry, and taphonomy (i.e., the study of processes that lead to fossilisation) suggest that nodules provide a mosaic of crevices at mm to cm scales, with stable hard surfaces, protected crevices, as well as gradients in porosity and chemistry. Scan electron microscopy showed that nodules are internally heterogeneous, and DM and SEM-EDX indicated variation in Mn, Fe, Co, Ni and Cu, reflecting alternating dense, coarse textures that created cracks and pores. Almost a quarter of nodule space was found to be porous (23.0 ± 6.83%) and larger nodules were found to have larger cracks. This complexity likely provided attachment points and refugia to fauna that were unavailable in the sediment. The connection between internal and surface cracks likely permits exchange of solutes and particulates allowing for colonisation from the sediment-water interface.

Microfossil evidence supported the view that these internal spaces are biologically available during growth as well as after initial crack formation, although initial crack formation, the timing and mode of incorporation cannot be resolved definitively. We observed abundant microfossils within nodules consistent with entry during nodule growth when laminae were still forming and permeable, and with later ingress through fractures followed by partial cementation. This would explain the mixture of forms and preservation states seen in the SEM images (Fig. 2). The presence of agglutinated foraminifera fragments and other biogenic particles within crevices indicated that at least some cavity structures were colonised by ancient organisms, although this does not necessarily demonstrate in-situ colonisation at the time of incorporation. The elevated Na and Cl contents may reflect retained or precipitated marine salts within the porous agglutinated tests and should not be interpreted as biological uptake. Although restricted to a single nodule, the result shows that incorporated biological material contributes to fine-scale chemical as well as structural heterogeneity within nodules. Sponge spicules were visible in the SEM images, and the presence of the Foraminifera species *Saccorhiza* sp., which uses sponge spicules to build its tests, indicates that nodules may have provided a habitat on the seabed for millions of years [22]. It has been proposed that nodule crevice Foraminifera represent primitive taxa that may have evolved at a slower rate than the sediment communities, but that their long-standing relationship with nodule-crevices is likely a result of the plasticity of their test structures, allowing them to mould their tests to any crevice large enough [22]. Together with the mapped networks of cracks analysed here, these observations indicate that nodules can host various fauna across growth stages.

This quantitative assessment of nodule crevice communities across multiple CCZ contract areas, including DNA-based comparisons of nodule-crevice and sediment meiofaunal assemblages, establishes a regional baseline for this poorly known component of abyssal biodiversity. Nodule interiors are rugose and structurally complex, a feature that likely contributes to the differences in meiofaunal assemblages between nodules and sediments in the CCZ [21,23,24]. Crevice fauna abundance and biomass increased with nodule volume; however, small nodules harboured a disproportionate amount of fauna, and the size distribution of nodules should therefore be considered when estimating the faunal abundance affected by nodule removal. Abundance of nematodes and copepods was significantly higher in the GSR compared to the BGR nodule exploration contract area, while foraminifera showed an inverse trend, pointing to taxa-specific community patterns. Molecular data showed that nodule samples were more similar to one another than to the nearby sediments, despite the ∼1000 km distance between sampling areas for copepods and an even greater distance for nematodes, underscoring that nodule-crevice assemblages were consistently distinct from sediment communities both morphologically [23,24] and at the 18S sequence-level (this study). Importantly, the effect of habitat was significant across dominant meiofaunal taxa and contributed at least 10 % to the community compositional difference for both nematodes and copepods, highlighting the potential importance of nodule crevices for the meiofaunal community composition of the CCZ benthos. Given the extended timescales that nodules take to form, preference for, or specialisation to, a nodule matrix would have been possible over evolutionary timescales. The high proportion of unclassified nematode ASVs and the divergent copepod lineages may reflect sequence divergence within the approximately 400-bp V1–V2 region of the 18S rRNA gene, possibly reflecting evolutionary shifts, but also the under-representation of deep-sea taxa in reference databases and constraints associated with taxonomic assignment. Importantly, the authors would like to note that the relatively low number of unique sequences recovered for both nematodes and copepods in the nodules relative to the sediments may have resulted from incomplete ethanol penetration into water-filled nodule crevices which may have reduced DNA quality and amplification success. The recovered sequences should therefore not be interpreted as a complete inventory of nodule-associated diversity. Nevertheless, nodule assemblages remained consistently distinct from sediment assemblages across locations and datasets. Because the same 18S V1–V2 primer pair was used throughout, this separation is unlikely to result from differences in marker choice.

The presence of nodules is thought to enhance diversity by providing a hard-substrate habitat to otherwise sediment-dominated abyssal plains [26,54], as seen in other deep-sea environments such as vents, seeps and seamounts, where habitat complexity promotes niche diversification [55,56]. Our results showed that within this broader context, individual nodules, whilst being considerably less porous than surrounding sediments and therefore containing less animals, hosted different meiobenthic communities to those of the sediments, alongside the macro- and megafaunal assemblages already documented on nodule surfaces [57,58]. Furthermore, the 244 nodules analysed here represent a small sample relative to the extent of nodule habitat in the CCZ, and current sampling remains insufficient to characterise the full diversity and distribution of nodule-crevice communities. Nodule collection will cause effectively permanent loss of the physical nodule habitat within the footprint of the mining activity, preventing recovery of nodule-associated communities at those locations on geological timescales. Whether this produces population- or species-level loss will depend on habitat specificity, regional distributions, connectivity and the amount and spatial configuration of nodule habitat retained elsewhere. These uncertainties should be explicitly considered in environmental management and conservation planning for the CCZ.

## Methods

### Sample collection

#### Nodules for habitat characterisation

Nodules used for DM were collected during three expeditions to the BGR and GSR exploration contract areas in the CCZ in 2021 (expedition GSRNOD21 aboard M/V *Normand Energy* [59], expedition Mangan2021 aboard M/V *Island Pride* [60]) and 2022 (expedition SO295 aboard R/V *Sonne* [61]). During the GSRNOD21 expedition, nodules were collected with the pre-prototype nodule collector vehicle “Patania II” (PATII) developed and deployed by GSR, as part of the nodule collection trial [17,59]. During the Mangan2021 expedition, nodules were sampled with the remotely operated vehicle ROV HD14 from Ocean Infinity (Austin, Texas, USA) and with a box corer [60], while during the SO295 expedition, nodules were sampled with push corers by ROV Kiel 6000 (Geomar, Kiel, Germany) [61]. Nodules for SEM were sampled with push corers and blade corers, that were all deployed by ROV Kiel 6000, and with a multi corer during the SO295 expedition. Additional nodules were sampled with a box corer and by ROV HD14 during the Mangan2021 expedition.

#### Nodules for crevice fauna analyses

Nodules used for the analysis of nodule crevice fauna were also collected in the BGR and GSR contract areas in 2019 (expeditions SO268-1 and SO268-2 aboard R/V *Sonne* [62]). Nodules were collected from the sediment surface of 16 box core samples [25] (Fig. 1) during the JPI Oceans Mining Impact II (MI2) project, specifically, in the BGR MI2 dredge site (i.e., a site in which a small-scale dredge experiment was conducted to evaluate the potential impacts of an upscale collector [16]). Samples were collected pre-impact to characterise baseline communities at the BGR MI2 trial site (i.e., the site where the PATII would be later deployed in 2021 during Mangan2021), and at the reference site (i.e., a nearby site containing undisturbed seafloor to be used as baseline control for comparison with disturbed areas). The same logic was applied to the GSR where samples were collected from the GSR MI2 trial site (pre-impact) and the GSR MI2 reference site.

Dimensions (length *L*, width *W*, and height *H*; all in mm) of all nodules were measured with a mechanical ruler [25] for all samples above 5 g. Samples below 5 g were not measured. Before preservation, visible epifauna were manually removed from each nodule, after which the nodules were thoroughly rinsed with 0.1 µm-filtered seawater following the approach of Pape et al. (2021) [24]. This step was intended to dislodge organisms remaining on the external nodule surface before the nodules were preserved to recover fauna retained within nodule crevices. The same sequence of processing steps was applied to all nodules and nodules were rinsed until sediment was no longer visibly washed off into the existing samples of the 0 – 3 cm sediment layer fraction of each boxcore. The first 25 nodules of each boxcore were fixed in 8% buffered formaldehyde for morphological analyses. All remaining nodules in the boxcore were split evenly. The first half was fixed in formaldehyde while the other half was fixed in 96% molecular-grade ethanol. Ethanol samples were stored at -20°C on board, until processing at NIOZ – Royal Netherlands Institute for Sea Research (Den Hoorn (Texel), Netherlands). Formaldehyde samples were stored at room temperature.

### Sample processing

#### Nodules processed for habitat characterisation

Internal structures and elemental composition of nodules were investigated at the Chair of Materials Sciences (LWK) of Paderborn University (Paderborn, Germany). Dimensions (length *L*, width *W*, and height *H*; all in mm) of all nodules were measured with a digital caliper. The ellipsoidal volume (*V_ellipsoid_*) of each nodule was calculated using the standard formula based on semi-axes:

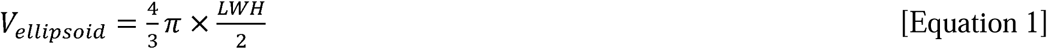

In order to ensure that an appropriate size range was used for downstream analysis, nodules were categorized by volume (ml) as ‘very small’ (0–10), ‘small’ (11–50), ‘medium’ (51–100), ‘large’ (101–200), and ‘very large’ (>200) [35]. To quantify the liveable space created by nodule crevices (i.e., dimensions of cracks and % porosity formed by dendritic growth structures), several nodules of each size class (*n*_very small_ = 5, *n*_small_ = 5, *n*_medium_ = 5, *n*_large_ = 8, *n*_very large_ = 1) were embedded individually in a two-component cold embedding medium (CEM1000 Blue Härter [article number: CEM055] + CEM 1000 Blue Pulver [article number: CEM033P]; Cloeren Technology GmbH, Wegberg, Germany). After hardening for 2 to 24 h depending on the nodule size, each nodule was divided with a bandsaw into two halves along its longitudinal axis. The cross sections were sanded with a double samples-polisher ‘Phoenix 2000’ (Jean Wirtz, Duesseldorf, Germany) using subsequently five different waterproof silicon carbide abrasive sheets (WS flex 18C; Hermes Schleifmittel GmbH, Hamburg, Germany) with grits of P80, P220, P500, P1200, and P2500.

The central section of each half nodule was scanned using the Keyence VHX-5000 DM (Keyence Corporation, Osaka, Japan) with a 100× universal zoom lens (Keyence VH-Z100UR, 100× to 1000×) and full ring lighting. As the size of a nodule section exceeded the observable image size of 20,000 (H) pixels × 20,000 (V) pixels when stitched by the microscope, several spatially overlapping scans were taken per nodule section. After the scans of the central section were completed, the two halves of nodules from the ‘large’ and ‘very large’ size classes were again divided by bandsaw, sanded, and one section per upper and lower halves (i.e., the so-called top and bottom sections) were scanned.

For a complete 2D digital image of the entire nodule section, the spatially overlapping scans were stitched together manually. Afterwards, the photomosaic was uploaded to ‘Keyence measurement software VHX-H2M2’ of the DM and the diameter (in µm) of internal and external cracks was measured as the distance between two points. When a crack was several cm long, multiple diameter measurements were taken. Information about the number and area (in µm^2^) of crevices, as well as the porosity of each nodule section were retrieved from the photomosaic with the automatic area measurement setting of the VHX-H2M2 software based on the brightness of the crevices and cracks.

To investigate the elemental composition of the nodule and the presence of microfossils inside the nodules, polished nodule sections and unpolished subsamples of nodules (size range: 12 mm 0.6 mm × 1.7 mm to 1.6 mm × 2.9 mm × 3.0 mm), that were broken off manually, were dried at 60 °C in a fan oven for 48 h. Afterwards, they were stored in a vacuum desiccator at 0.05 bar for several days. Dried, uncoated nodule samples were mounted on sample holders with copper tape and inserted into a Zeiss Ultra Plus field emission (FE) SEM (Carl Zeiss AG, Oberkochen, Germany). The polished samples were scanned under high vacuum conditions (10 – 5 mbar) with a focused electron beam produced by the Tungsten-zircon field-emission filament at an acceleration voltage of 20 kV. This allowed magnifications between 50× to 1,000×. The SEM measurements of unpolished nodule subsamples were performed under the same high vacuum pressure, but with acceleration voltages between 5 and 20 kV and magnifications ranging from 50× to 2,500×. The major element composition (in % weight) of structures was determined with the EDX detector Octane Pro (AMETEK, Berwyn, Pennsylvania, USA) at 20 kV acceleration voltage with 40,000 counts per second. Three 2D maps of major element composition of polished nodule sections (100× magnification) were created by SEM-EDX using an acceleration voltage of 20 kV and a dwell time of 200 µsec.

### Nodules processed for crevice fauna

To extract nodule crevice fauna, nodules fixed in both preservatives were frozen at -20°C for 24 h. This caused the otherwise hard substrate of the nodule to disintegrate into smaller sediment-adjacent sized particles, without observable damage or loss to biotic specimens across preservation methods (personal observation), a method developed at NIOZ. Afterwards, the disintegrated nodule samples were sieved through a 32 µm mesh size sieve to remove most of the fine particles whilst retaining the meiofauna. The material retained on the 32 µm sieve was centrifuged (6 min, 3,500 rpm) in a mixture of kaolin mud and levasil (LEVASIL®, Kurt Obermeier GmbH, Bad Berleburg, Germany) [63] to capture the meiofauna from the nodules in the supernatant. This procedure was repeated in total three times per sample to ensure that all meiofauna was extracted. Meiofauna was then sorted into higher taxa and counted per nodule under a Leica TL3000 Ergo stereo microscope (Leica Microsystems GmbH, Wetzlar, Germany) with a maximum magnification of 8×. The dominant meiofaunal Nematoda and Arthropoda (Copepoda) from nodules fixed in formaldehyde were mounted on microscopy slides for body size measurements. Specimens from samples fixed in ethanol were transferred back to ethanol until DNA extraction. Foraminifera were placed in cavity slides and viewed under a stereo zoom microscope. Foraminifera specimens were identified based on the taxonomic literature from the Pacific Ocean [47,64–66] and species names were cross-validated with the World Register of Marine Species (WoRMS [67]). Nematoda and Copepoda (96.7% of individuals) were photographed and measured in (version 1.52a) [68]. Nematode length *L* (including tail) and width *W* (in µm) were measured following Andrassy (1956) [69], and copepod length *L* and maximum width *W* (in µm) were measured following Warwick & Gee (1984) [70]. Based on these measurements, Nematoda dry mass *DM*_Nematoda_ (in µg) was calculated following Andrassy (1956) [69]:

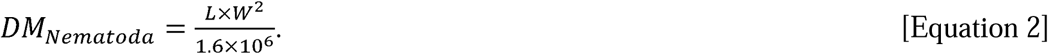

Copepoda *DM*_Copepoda_ (in µg) was calculated following Warwick & Gee (1984) [70]:

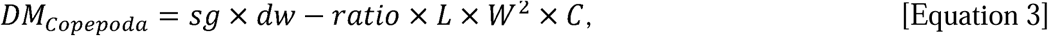

where *sg* is the specific gravity (1.13), *dw-ratio* is the dry mass/wet mass ratio (0.25), and *C* is the body-form conversion factor (i.e., semi-cylindrical Copepoda: 560, semi-cylindrical depressed Copepoda: 490 [70]).

### DNA extraction

Genomic DNA of Nematoda from nodule crevices (1 sample = 1 nodule community) was extracted as described in Macheriotou et al. (2020) [71] and amplified in triplicates. In short, the approximately 400-bp V1-V2 region of the 18S rRNA gene was amplified using SSU_F04 (GCTTGTCTCAAAGATTAAGCC) and SSU_R22 (GCCTGCTGCCTTCCTTGGA) primers [72]. The triplicate PCR products were run on a 1% agarose electrophoresis gel, before they were pooled, purified, and run on a tape station to confirm the length and size distribution of the PCR fragments. The pooled library was sequenced at Macrogen Europe BV (Amsterdam, Netherlands) on an Illumina MiSeq-v3 2 × 300 bp paired-end read run.

Genomic DNA of individual nodule crevice Copepoda was extracted using Chelex 100 Resin (Sigma Aldrich, Burlington, Massachusetts, United States) following the manufacturer’s protocol [73]. The V1-V2 hypervariable regions of the 18S rRNA gene were also amplified using the primers SSU_F04 and SSU_R22. Each sample was amplified in a 25 µl reaction containing 9.5 µl PCR-grade H_2_O, 12.5 µl AccuStart GelTrack PCR SuperMix (ThermoFisher Scientific, Waltham, Massachusetts, United States), 0.5 µl of each primer (10 pmol l ¹), and 2 µl of DNA template, using the following PCR conditions: 3 min at 94°C, 40 cycles of 30 s at 94°C, 45 s at 45°C, 45 s at 72°C, and afterwards 2 min at 72°C. PCR products were sequenced at Macrogen Europe (Amsterdam, Netherlands) using an ABI 3730XL Sanger sequencer.

### Data analysis

#### Nodules as habitat

Based on the major element composition of growth layer structures inside polymetallic nodules, they were classified as layer type 1, layer type 2.1, layer type 2.2, and layer type 3 [29]. Growth rate *R* (in mm My^-1^) of individual growth structures was calculated using two approaches. The first approach (Eq. 4, *R_Co_*) is based on the Co content of the growth structure [74] and the second approach (Eq. 5, *R_Fe+Mn_*) is based the Mn and Fe content of the structure [74]. These two approaches are complementary for deriving nodule age.

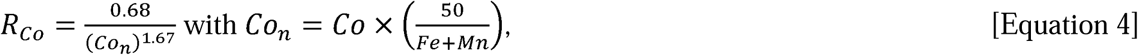

where Co, Fe, and Mn are reported in wt%.

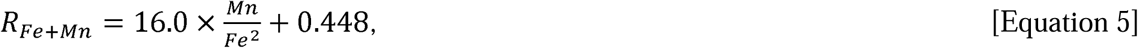

where Mn and Fe are reported in wt%.

#### Bioinformatics

Crevice meiofauna (this study) were compared with sediment infauna (published studies [16,17,71]) using 67 sediment samples from the BGR and GSR contract areas, which were collected in 2015 (included only in the PCoA [75] and Venn diagram) and 2019 (used for modelling and statistical comparison) (Sequence Read Archive: PRJNA889344). Metabarcoding pipelines were implemented using the *R* software (version 4.3.3) [76]. Specifically, for the Nematoda, published data were reanalysed together with nodule crevice data (Sequence Read Archive: PRJNA1331807) using the pipeline below.

Nematoda metabarcoding sequences were processed with the Dada2 pipeline [77] and classified using the Ribosomal Database Project (RDP) classifier [78] at 80% confidence. A combined eukaryotic reference database, including SILVA release 123 for QIIME1 [79] (99% OTUs) and an in-house database of Ghent University (Ghent, Belgium) with Sanger sequences of Nematoda (*n* = 18,991) [71], was first used. Amplicon Sequence Variants (ASVs) assigned to Nematoda were then extracted and reclassified using a second in-house database of Ghent University with Sanger sequences of marine Nematoda (*n* = 971) [16]. All genetic information along with metadata were stored in a *phyloseq* [80] object in *R* for downstream analysis. For the Nematoda, all ASVs (whether from nodules or sediments) with at least 5 reads in any sample were retained resulting in 67 unique ASVs from nodules and 8745 ASVs from sediments. All post-impact samples from the BGR were removed, leaving a purely baseline dataset. Community analysis was done on presence-absence by converting the OTU table to a binary matrix (all non-zero counts were set to 1), so that each ASV represents an occurrence in each sample. Given the size of the sediment dataset, stratified sampling was done to reduce the number of sequences to 200 for phylogeny construction, by down-sampling to an equal representation of rare, intermediate and abundant sediment lineages. All nodule sequences were retained. Nodule and sediment sequences were aligned using the MAFFT v7.490 plugin in Geneious Prime [80], the alignment was trimmed and cleaned in Geneious Prime, producing an alignment with length of 379 bp. A maximum-likelihood tree was built with IQ-TREE (1000 ultrafast bootstraps, BS) [81]. The tree was rooted on the most distant clade with a BS value of 100, and attached to the corresponding *phyloseq* object, yielding a nematode *phyloseq* object with presence–absence OTU table, sample metadata and 18S phylogeny. Species delimitation was performed with bPTP [82], based on phylogenetic branch lengths. Taxonomic classification was assigned from the RDP classifier.

From the nodule crevice Copepoda, 107 newly generated sequences were trimmed and assembled into contigs using Geneious Prime (version 2022.0.2; https://www.geneious.com). Ambiguous base calls were resolved and gaps removed where possible using the corresponding forward and reverse reads, and sequences with a high-quality score below 40% were excluded. Sediment metabarcoding ASVs and individually Sanger-sequenced nodule copepods, which targeted the same V1-V2 region of the 18S rRNA gene using the same primer pair, were then combined into a single dataset. Sanger sequences were imported from FASTA files and sample identifiers were parsed from the sequence headers. Because metabarcoding read counts and numbers of individually sequenced copepods are not quantitatively comparable, both datasets were converted to presence-absence before merging. All sequences were aligned and trimmed to the same homologous 309-bp region, then collapsed by exact nucleotide identity so that identical sequences were represented by a single unique 18S sequence variant, irrespective of sequencing method. Where multiple ASVs or Sanger sequences corresponded to the same variant, their occurrences were combined using a logical OR, such that the variant was recorded as present when detected at least once in a sample. This produced a single presence-absence matrix of unique copepod 18S sequence variants across nodule and sediment samples.

Phylogenetic reconstruction was performed as described for Nematoda. Species delimitation was performed with bPTP and taxonomic assignments and sequence percentage identities were obtained separately using BLAST [83] against the NCBI nucleotide database, with taxonomic information retrieved from the NCBI taxonomy database [84] using the *taxize* package [85].

### Statistical analysis

#### Relationships between nodule volume and structural characteristics of nodules

For all volume-based analyses, nodule ellipsoid volume was log_2_-transformed and mean-centred such that a unit increase represented a doubling in nodule volume. Relationships between ellipsoid nodule volume and structural characteristics were then analysed using mixed-effects models using the *ordinal* [86] and *glmmTMB* [87] package (version 1.1.14) in *R*, with section and contract area as fixed effects, and nodule or nodule-section identity as random intercepts where required. Model families were selected according to response type and fit: negative-binomial regression for overdispersed crevice counts, beta regression for proportional porosity data, cumulative-link regression for ordered median crevice-area values, and Student-t models for right-skewed mean crevice area which were log(1 + response)-transformed and crack diameters (ln-transformed). This response-specific approach follows previous deep-sea and polymetallic-nodule studies in which error distributions were matched to the statistical properties of ecological and environmental responses [15,31,88].

Additive, volume-by-section interaction and quadratic-volume models were compared using AICc [89], likelihood-ratio tests, and residual diagnostics. Final model adequacy was assessed using simulation-based residuals, and robustness was examined using central-section-only models, nodule-cluster bootstrapping, section-level median crack diameters, and leave-one-nodule-out analyses. Effects were reported with 95% CI and, where applicable, as percentage changes or ORs per doubling of nodule volume. Analyses were conducted using the R packages *tidyverse* [90], *glmmTMB* [87], *ordinal* [91], *DHARMa* [92], *performance* [93], *emmeans* [94], *broom.mixed* [95], and *MASS* [96].

#### Nodule crevice organisms

Abundance of crevice fauna per 10 cm^2^ was calculated by using the boxcore surface area (2,500 cm^2^) divided by the nodule surface area per boxcore. Diversity indices (species richness *S*, Shannon diversity index *H*′ [97], and Pielou’s evenness *J*′ [98]) were calculated for each nodule separately for Nematoda and Copepoda using the *vegan* package (version 2.6) [99], based on bPTP species delimitation results.

Relationships between nodule volume (log_2_-transformed and mean-centred) and faunal abundance, as well as biomass and size of dominant meiofauna (nematode length and copepod width in µm) were also analysed using mixed-effects models with the *glmmTMB* package [87], where models included exploration contract area (BGR and GSR) as a fixed effect and box core as a random intercept. Poisson and negative-binomial models with alternative dispersion structures were compared, and the simplest diagnostically acceptable model within two AICc units of the best-supported model was retained.

Combined Nematoda and Copepoda biomass was analysed using a Gamma mixed model. Log-transformed nematode length and copepod width were analysed using Student-t and Gaussian mixed models, respectively, with nodule and, where applicable, box core included as random effects. Model adequacy was assessed using simulation-based residual checks for departures from the expected residual distribution, over- or underdispersion, excess zeros and unexplained outliers. Differences between contract areas were tested by comparing models with and without contract area, and coefficients were reported as percentage changes per twofold increase in nodule volume. To test whether the greater representation of small nodules influenced the results, nodules were binned into the five size classes reported for the nodules used to characterise nodule habitat (very small-very large). Classes were repeatedly down-sampled to equal sample sizes while retaining composition between contract areas where possible. The selected models were refitted to 1,000 balanced datasets and compared with the complete-data estimates.

#### Nematoda

For Nematoda, community dissimilarities were quantified using Sørensen dissimilarities (binary Bray-Curtis) [100]. Because the phylogeny represented a subset of the full community, Sørensen dissimilarity was used for inferential comparisons.

To account for non-independence among samples collected within the same deployment, the nematode presence–absence ASV table was aggregated at the parent sampling-unit level, with nodule samples grouped by box core and sediment samples grouped by multi corer deployment. Within each sampling unit, an ASV was considered present when it occurred in at least one constituent molecular sample. This resulted in six independent nodule box cores and 16 independent sediment multi corer deployments. Habitat differences were tested using the model Sørensen dissimilarity ∼ contract area+ habitat, with marginal tests used to estimate the effect of each term while accounting for the other. Differences between contract areas were subsequently tested separately within nodule and sediment habitats. Community composition was first visualised using PCoA. Permutational multivariate analysis of variance (PERMANOVA) [101] with 9,999 permutations was used to test for differences in community composition among groups. To confirm that PERMANOVA results were not driven by heterogeneous within-group variability, homogeneity of multivariate dispersions was evaluated using the distance-to-centroid approach implemented using the betadisper function (9,999 permutations) in the *vegan* package.

#### Copepoda

For Copepoda, an 18S alignment and maximum-likelihood phylogeny spanning the complete set of sequence variants included in the community matrix (metabarcoding ASVs from sediments and Sanger haplotypes from nodules) was available. Therefore, Copepoda beta-diversity was quantified using unweighted UniFrac. For Copepoda, the presence-absence sequence-variant table was aggregated by an independent sampling unit, with nodule samples grouped into 13 box cores (*n*_BGR_ = 5, *n*_GSR_ = 8) and sediment samples grouped into 10 multi corer deployments, all from the GSR. Copepod beta-diversity was quantified using unweighted UniFrac because the maximum-likelihood phylogeny included the complete set of sequence variants in the community matrix. Habitat differences were tested using the model unweighted UniFrac ∼ contract area + habitat, with marginal PERMANOVA tests and 9,999 permutations. Contract-area differences were subsequently tested among nodule samples; the corresponding sediment comparison was not performed because independent BGR sediment parent units were unavailable. Homogeneity of multivariate dispersion was evaluated using betadisper with 9,999 permutations. Where differences between habitats were significant, SIMPER analyses were used as a driver analysis to quantify the contribution of individual ASVs/haplotypes to between-group differences. This analysis was restricted to the GSR, where independent nodule box cores and sediment multi corer deployments were available for both habitats. Because SIMPER cannot decompose UniFrac distances, sequence variants contributing to habitat dissimilarity were identified using presence-absence based on binary Bray-Curtis. SIMPER contributions from sequence variants assigned to the same genus were subsequently summed to obtain genus-level contributions. Size-based rarefaction and extrapolation were conducted on bPTP-delimited species derived from datasets containing 67 ASVs (Nematoda) and 107 Sanger sequences (Copepoda) and implemented through iNEXT (version 3.0.0) [102].

Whenever means are reported, data are presented as mean ± SD.

## References

1. Murray, J. & Renard, A.-F. Report on Deep-Sea Deposits Based on the Specimens Collected during the Voyage of H.M.S. Challenger in the Years 1872 to 1876. (HM Stationery Office, London, 1891).

2. International Seabed Authority. The geographical limits of the Clarion-Clipperton Zone management area, exploration areas, reserved areas and areas of particular environmental interest (APEIs) (ISBA/17/LTC/7, ISBA/18/C/22, ISBA/26/C/58, ISBA/26/C/43). Marine Gazetteer https://marineregions.org/gazetteer.php?p=details&id=64222 (2026).

3. Hein, J. R. Manganese nodules. Encyclopedia of Marine Geosciences 408–412 (2016).

4. Ku, T.-L. & Broecker, W. S. Uranium, thorium, and protactinium in a manganese nodule. Earth Planet. Sci. Lett. 2, 317–320 (1967).

5. Ku, T.-L. & Broecker, W. S. Radiochemical studies on manganese nodules of deep-sea origin. Deep Sea Research and Oceanographic Abstracts 16, 625–637 (1969).

6. Gollner, S. et al. Resilience of benthic deep-sea fauna to mining activities. Mar. Environ. Res. 129, 76–101 (2017).

7. Stratmann, T., Soetaert, K., Kersken, D. & Van Oevelen, D. Polymetallic nodules are essential for food-web integrity of a prospective deep-seabed mining area in Pacific abyssal plains. Sci. Rep. 11, 12238 (2021).

8. Rabone, M. et al. How many metazoan species live in the world’s largest mineral exploration region? Current Biology 33, 2383–2396.e5 (2023).

9. Washburn, T. W. et al. Patterns of Macrofaunal Biodiversity Across the Clarion-Clipperton Zone: An Area Targeted for Seabed Mining. Front. Mar. Sci. 8, 626571 (2021).

10. Stewart, E. C. D. et al. Biodiversity, biogeography, and connectivity of polychaetes in the world’s largest marine minerals exploration frontier. Divers. Distrib. 29, 727–747 (2023).

11. Jones, D. O. B. et al. Biological responses to disturbance from simulated deep-sea polymetallic nodule mining. PLoS One 12, e0171750 (2017).

12. Jones, D. O. B. et al. Long-term impact and biological recovery in a deep-sea mining track. Nature 642, 112–118 (2025).

13. Stewart, E. C. D. et al. Impacts of an industrial deep-sea mining trial on macrofaunal biodiversity. *Nat*. Ecol. Evol. 10, 318–329 (2026).

14. Stratmann, T. et al. Recovery of Holothuroidea population density, community composition and respiration activity after a deep-sea disturbance experiment. Limnol. Oceanogr. 63, 2140–2153 (2018).

15. Simon-Lledó, E. et al. Biological effects 26 years after simulated deep-sea mining. Sci. Rep. 9, 8040 (2019).

16. Lefaible, N. et al. Digging deep: lessons learned from meiofaunal responses to a disturbance experiment in the Clarion-Clipperton Zone. Marine Biodiversity 53, 48 (2023).

17. Lefaible, N. et al. Industrial mining trial for polymetallic nodules in the Clarion-Clipperton Zone indicates complex and variable disturbances of meiofaunal communities. Front. Mar. Sci. 11, 1380530 (2024).

18. Glover, A. G. et al. The environmental impacts of deep-sea mining. Current Biology 36, R400–R419 (2026).

19. Bussau, C. New deep-sea Tardigrada (Arthrotardigrada, Halechiniscidae) from a manganese nodule area of the eastern South Pacific. Zool. Scr. 21, 79–91 (1992).

20. Bussau, C. Taxonomische und ökologische Untersuchungen an Nematoden des Peru-Beckens. *PhD-Thesis* (Kiel University, 1993).

21. Thiel, H., Schriever, G., Bussau, C. & Borowski, C. Manganese nodule crevice fauna. Deep-Sea Research I 40, 419–423 (1993).

22. Maybury, C. A. Crevice Foraminifera from abyssal South East Pacific manganese nodules. in Microfossils and Oceanic Environments (eds. Moguilevsky, A. & Whatley, R.) 281–293 (University of Wales, Aberystwyth, 1996).

23. Singh, R., Sautya, S. & Ingole, B. S. The community structure of the deep-sea nematode community associated with polymetallic nodules in the Central Indian Ocean Basin. Deep-Sea Research II 161, 16–28 (2019).

24. Pape, E. et al. Potential impacts of polymetallic nodule removal on deep-sea meiofauna. Sci. Rep. 11, 19996 (2021).

25. Schoening, T. & Gazis, I.-Z. Sizes, weights and volumes of poly-metallic nodules from box cores taken during SONNE cruises SO268/1 and SO268/2. Dataset Preprint at 10.1594/PANGAEA.904962 (2019).

26. Kuhn, T., Wegorzewski, A. V., Rühlemann, C. & Vink, A. Composition, formation, and occurrence of polymetallic nodules. in Deep-Sea Mining (ed. Sharma, R.) 23–63 (Springer International Publishing, Cham, 2017). doi:10.1007/978-3-319-52557-0_2.

27. Halbach, P., Scherhag, C., Hebisch, U. & Marchig, V. Geochemical and mineralogical control of different genetic types of deep-sea nodules from the Pacific Ocean. Miner. Depos. 16, 59–84 (1981).

28. Hein, J. R. & Koschinsky, A. Deep-ocean ferromanganese crusts and nodules. in Treatise on Geochemistry (eds. Holland, H. & Turekian, K.) vol. 13 273–291 (Elsevier Ltd., Amsterdam, 2014).

29. Wegorzewski, A. V. & Kuhn, T. The influence of suboxic diagenesis on the formation of manganese nodules in the Clarion Clipperton nodule belt of the Pacific Ocean. Mar. Geol. 357, 123–138 (2014).

30. Blöthe, M. et al. Manganese-cycling microbial communities inside deep-sea manganese nodules. Environ. Sci. Technol. 49, 7692–7700 (2015).

31. Yapan, B. C., Janssen, F., Boetius, A., Haeckel, M. & Molari, M. Diversity and connectivity of bacterial communities in polymetallic nodule-rich abyssal plains (Eastern Tropical Pacific). Front. Mar. Sci. 13, 1759595 (2026).

32. Larock, P. A. & Ehrlich, H. L. Observations of bacterial microcolonies on the surface of ferromanganese nodules from Blake Plateau by scanning electron microscopy. Microb. Ecol. 2, 84–96 (1975).

33. Wang, X. et al. Distribution of microfossils within polymetallic nodules: Biogenic clusters within manganese layers. Marine Biotechnology 14, 96–105 (2012).

34. Jiang, X. D. et al. Characterization and Quantification of Magnetofossils within abyssal manganese nodules from the Western Pacific Ocean and implications for nodule formation. Geochemistry, Geophysics, Geosystems 21, 2019GC008811 (2020).

35. Stratmann, T. et al. Physical and chemical characteristics of deep-sea polymetallic nodules collected from the GSRNOD21, Mangan2021, and SO295 scientific expeditions to the CCZ. PANGAEA dataset Preprint at 10.1594/PANGAEA.996200 (2026).

36. Skowronek, A. et al. Chemostratigraphic and textural indicators of nucleation and growth of polymetallic nodules from the Clarion-Clipperton Fracture Zone (IOM claim area). Minerals 11, 868 (2021).

37. De Wilt, M., Diaz-Recio Lorenzo, C. & Gollner, S. Raw counts of meiofauna collected from crevices of polymetallic nodules from box cores taken during SONNE cruises SO268/1 and SO268/2. PANGAEA dataset Preprint at 10.1594/PANGAEA.992747 (2026).

38. Bussau, C., Schriever, G. & Thiel, H. Evaluation of abyssal metazoan meiofauna from a manganese nodule area of the eastern South Pacific. Vie et Milieu 45, 39–48 (1995).

39. Lefaible, N., Macheriotou, L., Vanreusel, A., Hauquier, F. & Pape, E. Meiofauna counts from multi corer samples from the Reference site within the Belgian License Area during SONNE cruises SO268/1 and SO268/2, Clarion-Clipperton Zone. PANGAEA dataset Preprint at 1594/PANGAEA.942156 (2022).

40. Lefaible, N. Meiofauna counts from multi corer samples from a dredge-experiment in the German License area during SONNE cruise SO268/2, Clarion-Clipperton Zone. PANGAEA dataset Preprint at 10.1594/PANGAEA.942174 (2022).

41. Clarke, K. R. Non-parametric multivariate analyses of changes in community structure. Australian Journal of Ecology 18, 117–143 (1993).

42. Baguley, J. G., Montagna, P. A., Hyde, L. J., Kalke, R. D. & Rowe, G. T. Metazoan meiofauna abundance in relation to environmental variables in the northern Gulf of Mexico deep sea. Deep-Sea Research I 53, 1344–1362 (2006).

43. Schmidt, C., Lins, L. & Brandt, A. Harpacticoida (Crustacea, Copepoda) across a longitudinal transect of the Vema Fracture Zone and along a depth gradient in the Puerto Rico trench. Deep-Sea Research II 148, 236–250 (2018).

44. Shulga, N., Abramov, S., Klyukina, A., Ryazantsev, K. & Gavrilov, S. Fast-growing Arctic Fe–Mn deposits from the Kara Sea as the refuges for cosmopolitan marine microorganisms. Sci. Rep. 12, 21967 (2022).

45. Tully, B. J. & Heidelberg, J. F. Microbial communities associated with ferromanganese nodules and the surrounding sediments. Front. Microbiol. 4, 1–10 (2013).

46. Lozupone, C. UniFrac: a new phylogenetic method for comparing microbial communities. Appl. Environ. Microbiol. 71, 8228 (2005).

47. Gooday, A. J. et al. The Biodiversity and Distribution of Abyssal Benthic Foraminifera and Their Possible Ecological Roles: A Synthesis Across the Clarion-Clipperton Zone. Frontiers in Marine Science vol. 8 Preprint at 10.3389/fmars.2021.634726 (2021).

48. Gooday, A. J. et al. Giant protists (Xenophyophores, Foraminifera) are exceptionally diverse in parts of the abyssal eastern Pacific licensed for polymetallic nodule exploration. Biol. Conserv. 207, 106–116 (2017).

49. Dudley, W. C. Cementation and iron concentration in foraminifera on manganese nodules. The Journal of Foraminiferal Research 6, 202–207 (1976).

50. Verlaan, P. A. Benthic recruitment and manganese crust formation on seamounts. Mar. Biol. 113, 171–174 (1992).

51. Verlaan, P. A. & Cronan, D. S. Origin and variability of resource-grade marine ferromanganese nodules and crusts in the Pacific Ocean: A review of biogeochemical and physical controls. Geochemistry 82, 125741 (2022).

52. Hedley, R. H. Cement and Iron in the Arenaceous Foraminifera. Micropaleontology 9, 433 (1963).

53. Heron Allen, E. & Earland, A. X. — On some Foraminifera from the North Sea, etc., dredged by the Fisheries Cruiser “Goldseeker” (International North Sea Investigations— Scotland). V. On Thurammina papillata Brady: a Study in Variation. Journal of the Royal Microscopical Society 37, 530–557 (1917).

54. Uhlenkott, K., Vink, A., Kuhn, T. & Martínez Arbizu, P. Predicting meiofauna abundance to define preservation and impact zones in a deep sea mining context using random forest modelling. Journal of Applied Ecology 57, 1210–1221 (2020).

55. Fontaneto, D. et al. Characteristics of meiofauna in extreme marine ecosystems: a review. Marine Biodiversity 48, 35–71 (2017).

56. Gollner, S. et al. Three-dimensional management needs of deep-sea hydrothermal vent ecosystems. Mar. Policy 185, 106959 (2026).

57. Veillette, J. et al. Ferromanganese nodule fauna in the tropical North Pacific Ocean: Species richness, faunal cover and spatial distribution. Deep-Sea Research I 54, 1912–1935 (2007).

58. Vanreusel, A., Hilário, A., Ribeiro, P. A., Menot, L. & Martínez Arbizu, P. Threatened by mining, polymetallic nodules are required to preserve abyssal epifauna. Sci. Rep. 6, 26808 (2016).

59. Peacock, T. The GSR Patania II Expedition: Technical Achievements & Scientific Learnings. (2023) doi:10.6084/m9.figshare.26397403.v1.

60. Vink, A. et al. MANGAN 2021 Cruise Report: Independent Scientific Monitoring of Two Collector Tests in the BGR and GSR Contract Areas for the Exploration of Polymetallic Nodules in the Equatorial NE Pacific. www.bgr.bund.de (2022) doi:10.25928/hw7d-fs42.

61. Haeckel, M., Janssen, F. & Martinez Arbizu, P. Cruise Report SO295: Assessing the Impacts of Polymetallic Nodule Mining on the Deep-Sea Environment after an Industrial Collector Test in the German and Belgian Contract Areas in the CCZ (31 October 2022 - 23 December 2022, Port Hueneme (USA) - Port Hueneme (USA)). (2023) doi:10.48433/cr_so295.

62. Haeckel, M. & Linke, P. Cruise Report SO268: Assessing the Impacts of Nodule Mining on the Deep-Sea Environment (NoduleMonitoring), Manzanillo (Mexico) = Vancover (Canada) 17.02. - 27.05.2019. www.geomar.de (2021) doi:10.3289/GEOMAR_REP_NS_59_2021.

63. Burgess, R. An improved protocol for separating meiofauna from sediments using colloidal silica sols. Mar. Ecol. Prog. Ser. 214, 161–165 (2001).

64. Brady, H. B. Report on the Foraminifera dredged by H. M. S. Challenger during the years 1873 - 1876Report on the Foraminifera dredged by H. M. S. Challenger during the years 1873 - 1876. in Reports of Science Research Voyage of H. M. S. Challenger. Zoology vols 9, 221–814 (London, 1884).

65. Goineau, A. & Gooday, A. J. Radiolarian tests as microhabitats for novel benthic foraminifera: Observations from the abyssal eastern Equatorial Pacific (Clarion-Clipperton Fracture Zone). Deep-Sea Research I 103, 73–85 (2015).

66. Gooday, A. J., Goineau, A. & Voltski, I. Abyssal foraminifera attached to polymetallic nodules from the eastern Clarion-Clipperton Fracture Zone: A preliminary description and comparison with North Atlantic dropstone assemblages. Marine Biodiversity 45, 391–412 (2015).

67. WoRMS Editorial Board. World Register of Marine Species. https://www.marinespecies.org (2025).

68. Abràmoff, M. D., Magalhães, P. J. & Ram, S. J. Image processing with ImageJ. Biophotonics International 11, 36–42 (2004).

69. Andrassy, I. Die Rauminhalts- und Gewichtsbestimmung der Fadenwurmer (Nematoden). Acta Zoologica 2, 1–15 (1956).

70. Warwick, R. M. & Gee, J. Community structure of estuarine meiobenthos. Mar. Ecol. Prog. Ser. 18, 97–111 (1984).

71. Macheriotou, L., Rigaux, A., Derycke, S. & Vanreusel, A. Phylogenetic clustering and rarity imply risk of local species extinction in prospective deep-sea mining areas of the Clarion–Clipperton Fracture Zone. Proceedings of the Royal Society B: Biological Sciences 287, (2020).

72. Blaxter, M. L. et al. A molecular evolutionary framework for the phylum Nematoda. Nature 392, 71–75 (1998).

73. Walsh, P. S., Metzger, D. A. & Higuchi, R. Chelex 100 as a Medium for Simple Extraction of DNA for PCR-Based Typing from Forensic Material. Biotechniques 54, 134–139 (2013).

74. Manheim, F. T. & Lane-Bostwick, C. M. Cobalt in ferromanganese crusts as a monitor of hydrothermal discharge on the Pacific sea floor. Nature 335, 59–62 (1988).

75. Gower, J. C. Some distance properties of latent root and vector methods used in multivariate analysis. Biometrika 53, 325–338 (1966).

76. R-Core Team. R: A language and environment for statistical computing. Preprint at https://www.r-project.org/ (2025).

77. Callahan, B. J. et al. DADA2: High-resolution sample inference from Illumina amplicon data. Nat. Methods 13, 581–583 (2016).

78. Cole, J. R. et al. Ribosomal Database Project: data and tools for high throughput rRNA analysis. Nucleic Acids Res. 42, D633–D642 (2014).

79. Quast, C. et al. The SILVA ribosomal RNA gene database project: Improved data processing and web-based tools. Nucleic Acids Res. 41, (2013).

80. Katoh, K. & Standley, D. M. MAFFT Multiple Sequence Alignment Software Version 7: Improvements in Performance and Usability. Mol. Biol. Evol. 30, 772–780 (2013).

81. Minh, B. Q. et al. IQ-TREE 2: New Models and Efficient Methods for Phylogenetic Inference in the Genomic Era. Mol. Biol. Evol. 37, 1530–1534 (2020).

82. Zhang, J., Kapli, P., Pavlidis, P. & Stamatakis, A. A general species delimitation method with applications to phylogenetic placements. Bioinformatics 29, 2869–2876 (2013).

83. Camacho, C. et al. BLAST+: architecture and applications. BMC Bioinformatics 10, 421 (2009).

84. Wheeler, D. L. Database resources of the National Center for Biotechnology. Nucleic Acids Res. 31, 28–33 (2003).

85. Chamberlain, S. A. & Szöcs, E. taxize: taxonomic search and retrieval in R. F1000Res. 2, 191 (2013).

86. McCullagh, P. Regression Models for Ordinal Data. J. R. Stat. Soc. Series B Stat. Methodol. 42, 109–127 (1980).

87. Brooks, M. E. et al. glmmTMB Balances Speed and Flexibility Among Packages for Zero-inflated Generalized Linear Mixed Modeling. R J. 9, 378 (2017).

88. Gaikwad, S. et al. Macrobenthic communities in the polymetallic nodule field, Indian Ocean, based on multicore and box core analysis. Front. Mar. Sci. 11, 2024.1395892 (2024).

89. Hurvich, C. M. & Tsai, C.-L. Regression and time series model selection in small samples. Biometrika 76, 297–307 (1989).

90. Wickham, H. et al. Welcome to the Tidyverse. J. Open Source Softw. 4, 1686 (2019).

91. Christensen, R. H. B. ordinal: Regression Models for Ordinal Data. Preprint at (2026).

92. Hartig, F. & Hartig, M. F. Package ‘dharma’. Preprint at (2017).

93. Lüdecke, D., Ben-Shachar, M., Patil, I., Waggoner, P. & Makowski, D. performance: An R Package for Assessment, Comparison and Testing of Statistical Models. J. Open Source Softw. 6, 3139 (2021).

94. Lenth, R. emmeans: Estimated Marginal Means, aka Least-Squares Means. Preprint at (2023).

95. Bolker, B., Robinson, D., Menne, D., Gabry, J. & Buerkner, P. Package ‘broom. mixed’. Comprehensive R Archive Network.

96. Venables, W. N. & Ripley, B. D. *Modern Applied Statistics with S*. (Springer Science & Business Media, 2013).

97. Shannon, C. E. A mathematical theory of communication. The Bell system technical journal 27, 379–423 (1948).

98. Pielou, E. C. The measurement of diversity in different types of biological collections. J. Theor. Biol. 13, 131–144 (1966).

99. Oksanen, J., et al. vegan: Community ecology package. Preprint at https://cran.r-project.org/package=vegan. (2017).

100. Bray, J. R. & Curtis, J. T. An ordination of the upland forest communities of southern Wisconsin. Ecol. Monogr. 27, 325–349 (1957).

101. Anderson, M. J. Permutational Multivariate Analysis of Variance (PERMANOVA). in Wiley StatsRef: Statistics Reference Online 1–15 (Wiley, 2017). doi:10.1002/9781118445112.stat07841.

102. Hsieh, T. C., Ma, K. H. & Chao, A. iNEXT: An R package for rarefaction and extrapolation of species diversity (Hill numbers). Methods Ecol. Evol. 7, 1451–1456 (2016).

